## Supplemental Tables for "BINSEQ: A Family of High-Performance Binary Formats for Nucleotide Sequences"

### 1 Supplementary Materials and Data

#### 1.1 Format Specifications

##### 1.1.1 Shared Specification

*Table 4: Nucleotide Encoding. Standard nucleotides use two-bit encoding. In four-bit mode, these values are preserved in the least significant bits with leading zeros, while N requires the full four-bit representation.*

| Nucleotide | two-bit | four-bit |
|---|---|---|
| A | 00 | 0000 |
| C | 01 | 0001 |
| G | 10 | 0010 |
| T | 11 | 0011 |
| N | — | 1111 |

### 1.1.2 BQ

*Table 5: BQ Header Structure (32 bytes)*

| Offset | Size (bytes) | Field | Type | Description |
|---|---|---|---|---|
| 0 | 4 | magic | uint32 | Magic number (0x42534551) |
| 4 | 1 | version | uint8 | Format version (currently 2) |
| 5 | 4 | slen | uint32 | Sequence length (primary) |
| 9 | 4 | xlen | uint32 | Sequence length (secondary) |
| 13 | 1 | bits | uint8 | Number of bits per nucleotide (2, 4) |
| 14 | 1 | flags | bool | Records are prefixed by a flag uint64 |
| 15 | 17 | reserved | [uint8] | Reserved for future extensions |

##### 1.1.3 VBQ

*Table 6: VBQ Header Structure (32 bytes)*

| Offset | Size (bytes) | Field | Type | Description |
|---|---|---|---|---|
| 0 | 4 | magic | uint32 | Magic number (0x51455356) |
| 4 | 1 | format | uint8 | Format version (currently 1) |
| 5 | 8 | block | uint64 | Virtual size of all blocks in bytes |
| 13 | 1 | qual | bool | Records include quality scores |
| 14 | 1 | compressed | bool | Blocks are compressed |
| 15 | 1 | paired | bool | Records consist of sequence pairs |
| 16 | 1 | bits | uint8 | Number of bits per nucleotide (2, 4) |
| 17 | 1 | headers | bool | Records include sequence headers |
| 18 | 1 | flags | bool | Records are prefixed by a flag uint64 |
| 19 | 13 | reserved | [uint8] | Reserved bytes for future extensions |

*Table 7: VBQ Block Header (32 bytes)*

| Offset | Size (bytes) | Field | Type | Description |
|---|---|---|---|---|
| 0 | 8 | magic | uint64 | Magic number (0x5145534B434F4C42) |
| 8 | 8 | size | uint64 | True size of record block (can vary if compressed) |
| 16 | 4 | records | uint32 | Number of records in block |
| 20 | 12 | reserved | [uint8] | Reserved bytes for future extensions |

*Table 8: VBQ Record. B represents the number of bits per word and is 32 for two-bit encodings or 16 for four-bit.*

| **Field** | **Type** | **Size (bytes)** | **Description** |
|---|---|---|---|
| flag | uint64 | 8 | Binary flag for the record |
| slen | uint64 | 8 | Primary sequence length |
| xlen | uint64 | 8 | Extended sequence length |
| sbuf | [uint64] | $\lceil \frac{\text{slen}}{B} \rceil$ | Encoded primary sequence |
| squal | [uint8] | slen if paired else 0 | Primary sequence quality scores |
| sheader | [uint8] | 8 + Variable | Sequence Header. The first uint64 (8 bytes) defines the length in bytes of the header. The remaining bytes contain the header data. |
| xbuf | [uint64] | $\lceil \frac{\text{xlen}}{B} \rceil$ | Encoded extended sequence |
| xqual | [uint8] | xlen if paired else 0 | Extended sequence quality scores |
| xheader | [uint8] | 8 + Variable | Sequence Header. The first uint64 (8 bytes) defines the length in bytes of the header. The remaining bytes contain the header data. |

*Table 9: VBQ Index Header (32 bytes)*

| **Offset** | **Size (bytes)** | **Field** | **Type** | **Description** |
|---|---|---|---|---|
| 0 | 8 | magic | uint64 | Magic number (0x5845444e49514256) |
| 8 | 8 | bytes | uint64 | Number of bytes in the file pair) |
| 16 | 16 | reserved | [uint8] | Reserved bytes for future extensions |

*Table 10: VBQ Index Range (32 bytes)*

| **Offset** | **Size (bytes)** | **Field** | **Type** | **Description** |
|---|---|---|---|---|
| 0 | 8 | offset | uint64 | File offset of block start (bytes) |
| 8 | 8 | len | uint64 | Size of the block (bytes) |
| 16 | 4 | records | uint64 | Number of records in block |
| 20 | 4 | cumulative | uint64 | Number of records before block |
| 24 | 8 | reserved | [uint8] | Reserved bytes for future extensions |

#### 1.2 Processing

*Table 11: Parallel Processing Hooks*

| Hook | Type | Description |
|---|---|---|
| *process_record* | map | Process a single record (primary / extended) |
| *on_batch_complete* | reduce | Called when a batch of records has completed processing |
